# SFS_CODE: More Efficient and Flexible Forward Simulations

**DOI:** 10.1101/025064

**Authors:** Ryan D. Hernandez, Lawrence H. Uricchio

## Abstract

**SUMMARY:** Modern implementations of forward population genetic simulations are efficient and flexible, enabling the exploration of complex models that may otherwise be intractable. Here we describe an updated version of `SFS_CODE`, which has increased efficiency and includes many novel features. Among these features is an arbitrary model of dominance, the ability to simulate partial and soft selective sweeps, as well as track the trajectories of mutations and/or ancestries across multiple populations under complex models that are not possible under a coalescent framework. We also release `sfs_coder`, a Python wrapper to `SFS_CODE` allowing the user to easily generate command lines for common models of demography, selection, and human genome structure, as well as parse and simulate phenotypes from `SFS_CODE` output.

**Availability and Implementation:** Our open source software is written in `C` and Python, and are available under the GNU General Public License at http://sfscode.sourceforge.net.

**Supplementary information:** Detailed usage information is available from the project website at http://sfscode.sourceforge.net.

## 1 INTRODUCTION AND FEATURES

Efficient, highly flexible forward genetic simulations are essential components of several lines of research. In population and evolutionary genetics, they are necessary to evaluate the robustness of theoretical models (Bank *et al.* 2014); in association studies, they are crucial for measuring statistical power (Uricchio *et al.* 2015); and future applications may include statistical inference (Buzbas & Rosenberg 2015).

`SFS_CODE` is a computationally efficient implementation of a population genetic forward simulation. It has been used to understand the evolutionary forces driving patterns of genetic variation across many species (e.g. humans, Drosophila, plants, and coral), characterize complex demographic models, jointly model the effect of positive and negative selection, as well as designing statistical methods and sequencing studies. Since its introduction, `SFS_CODE` has undergone extensive revisions, and continues to be among the most efficient and flexible simulators available. Table 1 highlights novel features. In particular, efficient conditional simulations are now possible (e.g., using `--trackTrajectory` to retain simulations with specific terminal conditions, or `--mutation` to follow a specified trajectory).

We also introduced `sfs_coder`, a Python-based `SFS_CODE` interface that easily generates command-lines for several models of human demography, selection, and genome structure, as well as parses and analyzes `SFS_CODE` output, simulates selection-based phenotypes, and provides rescaling methods for simulations of linked positive selection.

**Table 1.** Key novel features added to `SFS_CODE`

| Option | Description |
| --- | --- |
| `--sampSize` | Sample populations at defined times |
| `--selDistType` | Dominance, selection on standing variation |
| `--rateClassLoci` | Locus-specific mutation rates |
| `--admix` | Join multiple populations at specified time |
| `--trackAncestry` | Report ancestry of each locus |
| `--trackTrajectory` | Follow frequency trajectory of an allele |
| `--mutation` | Introduce mutations; follow specified trajectory |

See documentation for details

## 2 METHODS

Efficiency gains in `SFS_CODE` have been achieved by storing all unique haplotypes in a locus as a collection of mutations in a randomized splay tree (a binary search tree that self-organizes upon accessing an element with probability *p*; (Albers & Karpinski 2002)). Individuals in a population are then stored as arrays of haplotypes across loci. Since many haplotypes are introduced and lost each generation, `SFS_CODE` efficiently reuses lost haplotypes to reduce allocation and deallocation overhead.

While highly efficient for many use cases, these data structures can come at a cost when the number of unique haplotypes is large relative to the population size (e.g., long locus length or high mutation/recombination rate). `SFS_CODE` therefore benefits if long chromosomes are partitioned into smaller linked segments. The ideal partitioning depends on several parameters, and can be optimized heuristically with simulations that have a short burn-in time using the Perl script `optimizeLL.pl` now included in the distribution.

### 2.1 Comparing performance

We compared the run time of simulations using `SFS_CODE` to several other efficient forward simulation programs that have recently been released: `FFPopSim` (Zanini & Neher 2012), `fwdpp_ind` (Thornton 2014), and `SLiM` (Messer 2013). Figure 1 shows the results, with the coalescent simulator `ms` (Hudson 2002) as a reference (which are often below scale as plotted). We varied the population size (*N*) and total locus length (*L*) for two values of the mutation (θ) and recombination (ρ) rates (roughly corresponding to human and Drosophila parameters). We find that for short loci, `FFPopSim` can be extremely efficient, but it does not scale well with *L*. `SFS_CODE` (and the locus-optimized version) is often the most efficient. For large *L* with unrealistically small *N*, forward simulations can be faster than the coalescent simulator `ms` (which has ∼exponential runtime in *L* with recombination; though see (Marjoram & Wall 2006)). All simulations were run on an Intel Xeon 2.66GHz CPU.

**Fig. 1.**
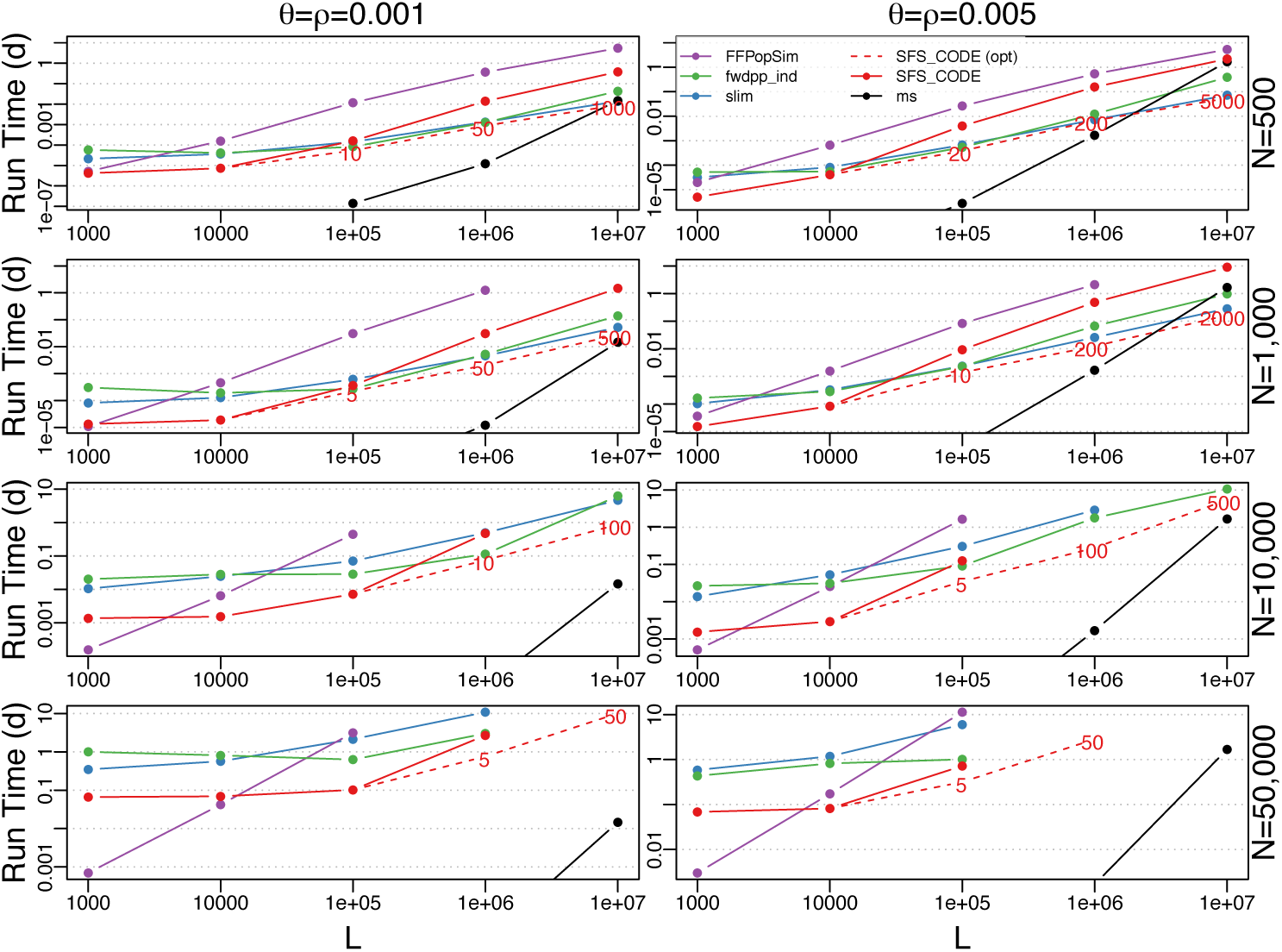
Mean run times across simulators, representing 12.55 CPU years of computation (10 iterations per point). Each panel is a function of simulated sequence length, with lower (human; left) and higher (Drosophila; right) mutation (θ) and recombination (ρ) rates. The coalescent simulator `ms` is included in black for reference. Dashed red lines are a locus-optimized version of `SFS_CODE`, with the optimal number of loci indicated in graph. Missing points represent parameter combinations that did not complete within 14 days.

## ACKNOWLEDGEMENTS

*Funding*: This work was supported by a grant from the National Institutes of Health (1R01HG007644) and a Sloan Foundation Research Fellowship to RDH.

